# *De novo* Protein Sequence Design Based on Deep Learning and Validation on CalB Hydrolase

**DOI:** 10.1101/2023.08.01.551444

**Authors:** Junxi Mu, Zhenxin Li, Bo Zhang, Qi Zhang, Jamshed Iqbal, Abdul Wadood, Ting Wei, Yan Feng, Haifeng Chen

**Affiliations:** Department of Bioinformatics and Biostatistics, School of Life Sciences and Biotechnology, Shanghai Jiao Tong University, 800 Dongchuan Road, Shanghai, 200240, China; State Key Laboratory of Microbial Metabolism, School of Life Sciences and Biotechnology, and Joint International Research Laboratory of Metabolic Developmental Sciences, Shanghai Jiao Tong University, 800 Dongchuan Road, Shanghai, 200240, China; Center for Life Sciences, Academy for Advanced Interdisciplinary Studies, Peking University, 5 Yiheyuan Road, Beijing, 100871, China.; Centre for Advanced Drug Research, COMSATS University Islamabad, Abbottabad Campus, Abbottabad, 22060, Pakistan; Department of Biochemistry, Abdul Wali Khan University Mardan, Mardan, 23200, Pakistan

**Keywords:** *De novo* Protein Design, Functional Verification, CalB, Artificial Intelligence

## Abstract

Protein design is central to nearly all protein engineering problems, as it can enable the creation of proteins with new biological function, such as improving the catalytic efficiency of enzymes. As one of the key tasks of protein design, fixed-backbone protein sequence design aims to design novel sequence that would fold into a given protein backbone structure. However, current sequence design methods have limitations in terms of low sequence diversity and experimental validation of designed protein function, which cannot meet the needs of functional protein design. We firstly constructed Graphormer-based Protein Design (GPD) model that directly applies Transformer to graph-based representation of 3D protein structure, and added Gaussian noise and sequence random mask to node features to improve the sequence recovery and diversity. Additionally, functional filtering based on the structure folding, solubility, and function were performed to improve the success rate in experiments. The process of “sequence design-functional filtering -functional experiment” was carried out for CalB hydrolase. The experimental results showed that the specify activity of designed protein improved 1.7 times than CalB wild type. This design and filtering platform will be a valuable tool for generating industrial enzymes and protein drugs with specific functions.

## 1 Introduction

Protein design is central to nearly all protein engineering problems and has a wide range of applications in enzyme engineers (designed enzymes with robust and active catalytic efficiency) [1] and therapeutics (designed immune proteins with augmented therapeutic affinity) [2]. *De novo* protein design aims to design novel amino acid that encode proteins with desired properties[3]. *De novo* Protein design includes two key tasks: protein backbone design and sequence design. Protein sequence design is also called the inverse protein folding problem. Here we focus on fixed-backbone protein sequence design. The objective of protein sequence design is to design a sequence that would fold into a given protein backbone structure. Specifically, the designed sequences not only need to fold into the desired structure, but also have the specified functions or required properties [4].

A number of works have been developed for protein sequence design. The main approaches for fixed-backbone protein sequence design can be divided into two categories: *Classical physical principle-based protein sequence design* and *deep learning-based protein sequence design* [5, 6]. Classical physical principle-based protein design, such as the leading protein design framework Rosetta [7], seeks to minimize the parametric energy function of the target structure by searching for the combination of sequence and conformations [3]. Classical physical principle-based approaches depend both on the accuracy of energy functions for protein physics and also on the efficiency of sampling algorithms, which the accuracy and computational speed need to be further improved [8].

With the rapid development of deep learning, recently deep learning-based protein design has shown the promising results to generate protein sequences that autonomously fold into a given protein backbone. Deep learning-based protein sequence design could provide fast and precise protein design, and has also caused a revolution in the field of computational protein design[8]. Supplementary Fig. 1 showed all the deep learning-based protein sequence design methods[9–21]. Only a few methods so far have reported experimental examinations of designed protein sequences: 3DCNN[15], ABACUS-R[16], and ProteinMPNN[18] designed sequences were experimental examinations by crystallograph; ProteinSolver[13] and ProDESIGN-LE[21] designed sequences were shown to have desired secondary structure contents according to circular dichroism signatures.

These methods have limitations in terms of protein functional validation and low sequence diversity, which cannot meet the needs of functional protein design. only a few methods have reported experimental structures of designed sequences[13, 15, 16, 18, 21], none of methods analyzed the protein function of designed sequences. Ideally, the designed sequences should have better performance than wild-type. Additionally, existed methods only sought to improve the sequence recovery between designed sequences and corresponding native sequence, and had limited the ability to explore the diversity between designed sequences, leading to designed sequences are overly monotonous and lack the diversity and variability. It is necessary to expand the sequence space landscape, and improve the diversity and variability of designed sequences.

Here, we introduced Graphormer-based Protein Design (GPD) toolbox, that inspired by Graphormer to directly apply Transformer to graph-based representation of 3D protein structure for protein sequence design and added a normally distributed random matrix in node features to increase the diversity of designed sequences. Additionally, functional filtering based on the structure folding, solubility, and function were performed to improve the success rate in experiments. We used GPD to design 1,000,000 de novo sequences of CalB hydrolase. Nine sequences after functional filtering were examined by wet experiments. The solubility of 9 designed sequences was 55.6% and the specify activity of one designed sequence was higher than CalB wild type. Graphormer-based Protein Design (GPD) toolbox was publicly available (https://yu.life.sjtu.edu.cn/ChenLab/GPDGenerator/). This web server allows the user automatically to generate protein sequence given the 3D protein structure and to fix the amino acids of interest at specified positions.

## 2 Results

### 2.1 The GPD architecture

Graphormer-based Protein Design (GPD) model directly applies Transformer to graph-based representation of 3D protein structure and adds a normally distributed random matrix in node features to increase the diversity of protein designed sequences. The node features contained the main-chain dihedral angle, the secondary classification, the centrality of each residue, the pre-designed protein sequence, and a tensor of random seed. The edge features contained the distances, the movement vectors, the shortest pathway, and the rotation quaternions.

### 2.2 Performance of GPD model

The CATH 40% sequential non-redundancy dataset has been used for GPD training. The split radio between training, validating and testing set was 29868:1000:103. We additionally test the performance of GPD using 39 *de novo* proteins, and 14 *de novo* proteins, which have meaningful structural differences from proteins belonging to natural folds [22, 23]. We compared the GPD with the widely used design approaches, including ProteinSolver, Structure Transformer, ESM-IF1, and ProteinMPNN. We used four evaluation criteria to systematically evaluate the performance of different methods from sequence and structure levels.

**Recovery**. The proportion of the same amino acids at equivalent position between the native sequence and the designed sequence.

**Diversity**. Diversity between designed sequences. Ideally, the designed sequences should cover a wide range of sequence space landscape and have higher diversity.

**Predicted Local Distance Difference Test scores (pLDDT)**. pLDDT of the ESMFold predicted structures.

**Root-Mean-Square Deviation (RMSD)**. Aligning the ESMFold predicted structures with the corresponding native structures .

#### 2.2.1 Designed sequences compared with native sequences

For GPD model, the average recovery between the designed sequences and corresponding native sequence is 46.2%±5.1%, 31.8%±5.8%, and 27.9%±5.4% on 14 *de novo* protein (Fig. 2a), 39 *de novo* proteins (Supplementary Fig. 2), and 103 single chains proteins (Supplementary Fig. 3), respectively. The average diversity between designed sequence is 21.9%±2.4% (Fig. 2b), 25.1%±3.3% (Supplementary Fig. 2), and 28%±5.6% (Supplementary Fig. 3). The recovery and diversity of our GDP model is significant higher than ProteinMPNN (P value=7.97e-3 and 1.80e-10) on single chain proteins. The recovery of ProteinMPNN is higher than GPD model on 14 *de novo* protein (P value=0.02) and 39 *de novo* proteins (P value=3.78e-05), However, the diversity of GPD is higher than ProteinMPNN on 14 *de novo* protein (P value=1.71e-03) and 39 *de novo* proteins (P value=2.55e-11).

**Fig. 1.**
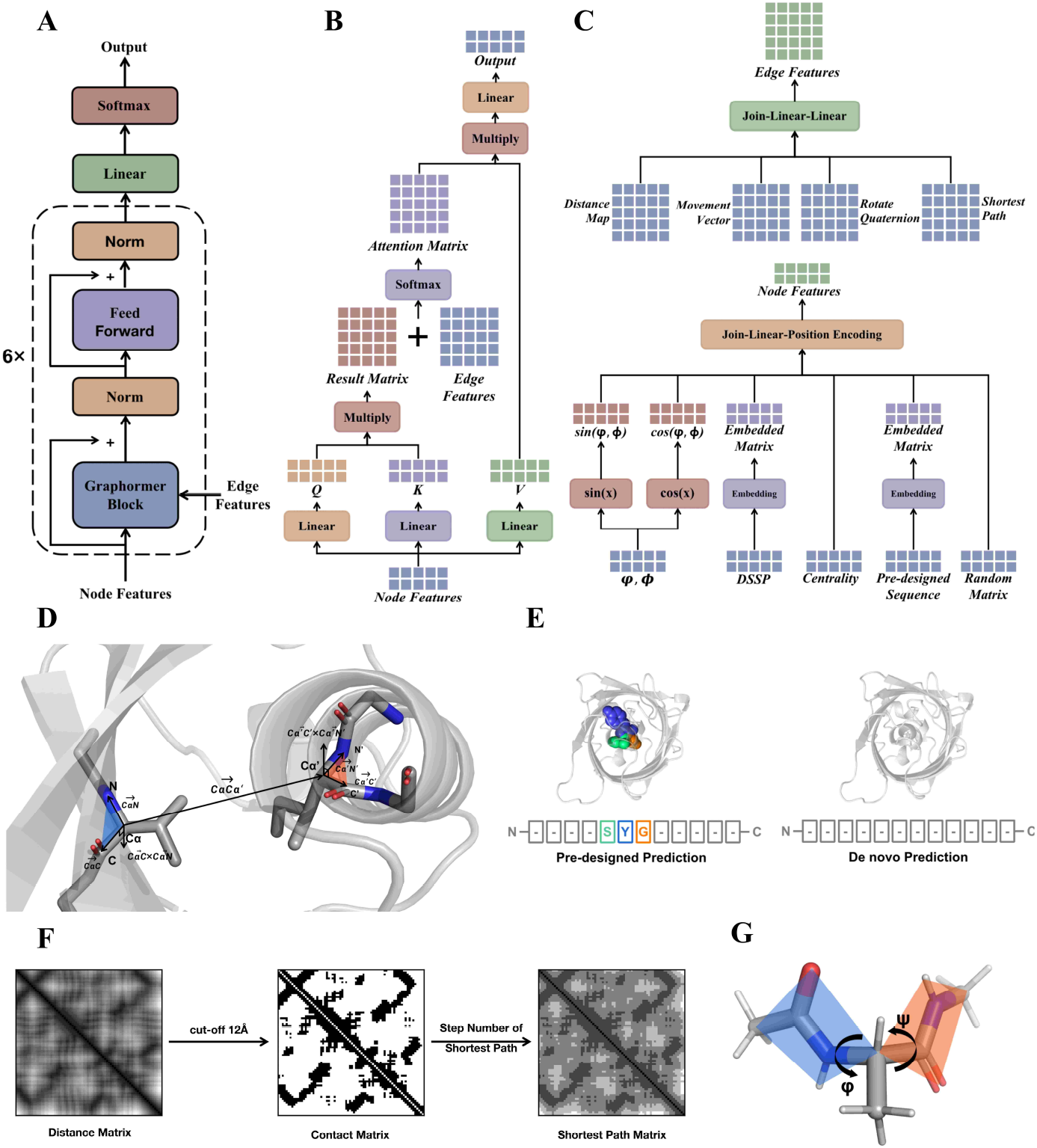
The GPD architecture. **a**, The overall architecture of GPD. **b,** The architecture of the Graphormer block. **c,** The embedding process of edge features and node features. **d,** The calculation process of distance map, local movement vector, and rotate quaternion. **e,** Two different ways of sequence prediction. **f,** The calculation of the shortest pathway matrix. **g,** The dihedral angles of residual backbone.

**Fig. 2.**
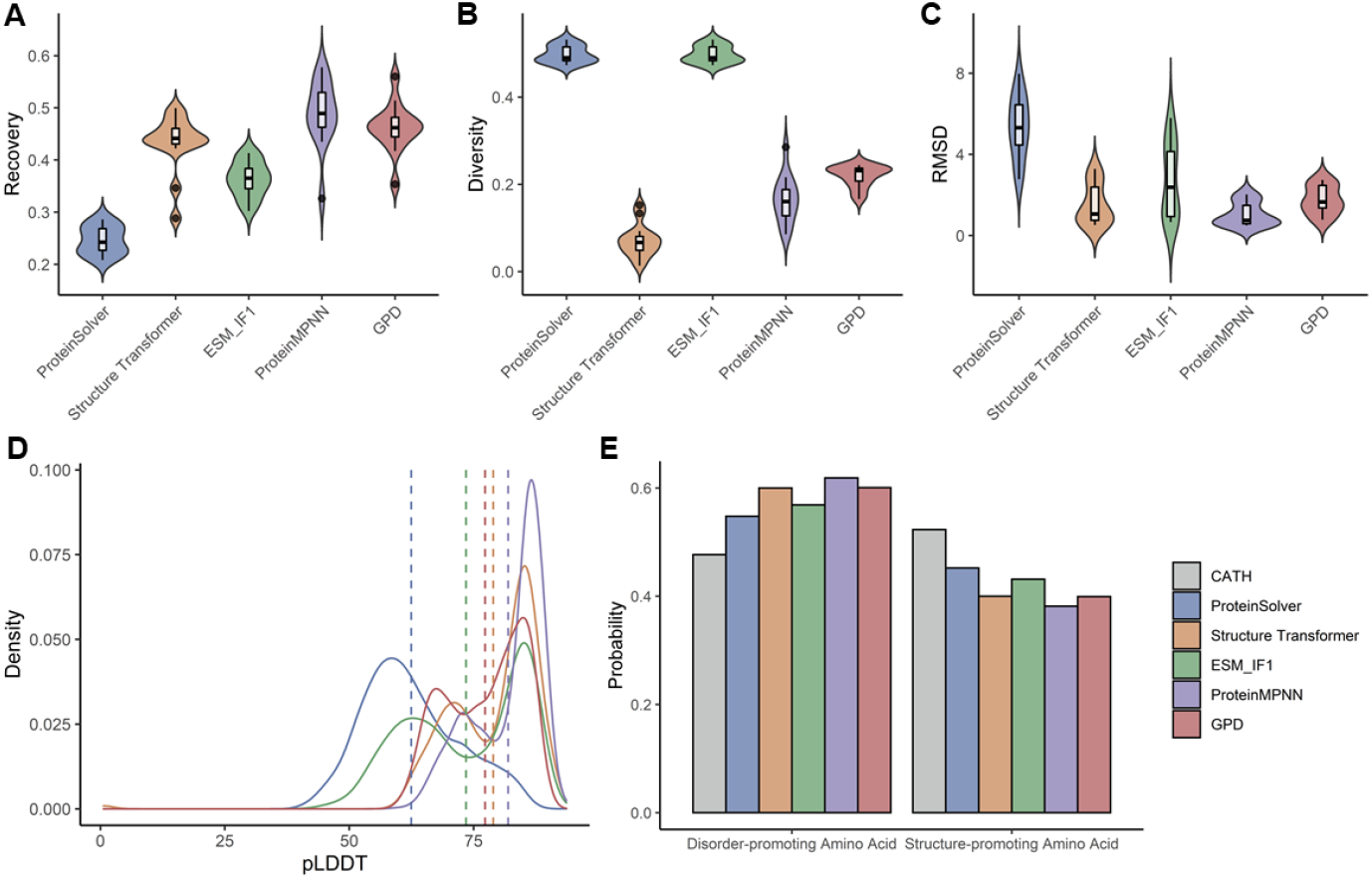
Comparison of designed sequence for five methods on 14 *de novo* proteins. **a**, the sequence recovery between the designed sequence and the native sequence of the target structure. **b,** The diversity of designed sequences. **c,** RMSD for aligning the ESMFold predicted structures with the corresponding native structures. **d,** The pLDDT scores of the ESMFold predicted structures. **e,** The frequency of disorder-promoting amino acids (alanine, glycine, proline, arginine, glutamine, serine, glutamic acid, and lysine) and structure-promoting amino acids (other twelve residues).

Recovery and Diversity are two metrics in fixed-backbone sequence design, and affect each other. Higher recovery would sacrifice the sequence diversity, and vice versa. It is necessary to consider recovery and diversity at the same time. Two proteins with greater than 35% sequence recovery are quite likely to have similar structures and perform similar functions[24]. For de novo proteins, the recovery is higher than 30% for all five methods (except Proteinsolver), and it is necessary to expand the sequence space landscape and improve the variability of designed sequences (Table A1). The higher diversity means the designed sequences have more diversity and cover a wide range of sequence space landscape. our GPD model could balance the recovery and diversity.

#### 2.2.2 Structure prediction on designed sequences by ESMFold

We designed 100 sequences for each protein using all five methods, and then applied ESMFold to predict the structures of 10300 designed sequences for 103 single chain proteins and 5300 designed sequences for 53 *de novo* proteins. The mean RMSD between the EMSFold predicted structures and the corresponding native structures, and the density plots of the pLDDT scores were shown in Fig. 2c and Fig. 2d (14 *de novo* protein), Supplementary Fig. 2 (39 *de novo* proteins), and Supplementary Fig. 3 (single chains proteins). The mean RMSD is 1.754^°^A, 3.647^°^A, and 7.303^°^A on 14 *de novo* protein (Fig. 2c), 39 *de novo* proteins (Supplementary Fig. 2), and single chains proteins (Supplementary Fig. 3), respectively. The percentage of designed sequences with RMSD less than 2^°^A is 68.29%, 39.51%, and 5.84%. All methods has the better performance on 14 *de novo* protein (Table A1). Supplementary Fig. 4 showed the predicted structure with the minimum RMSD. The ProteinMPNN designed sequences have the higher protein folding ability than other methods, even so, the percentage of designed sequences with RMSD less than 2^°^A is only 23.6% for single chain proteins. All these methods still have the deficiencies of protein folding, and this will lead to experimental failure. The protein folding ability is the important metrics to evaluate the performance of different methods.

Based on the folding ability, amino acids can be classified into two categories: disorder-promoting amino acids (alanine, glycine, proline, arginine, glutamine, serine, glutamic acid, and lysine) and structure-promoting amino acids (other twelve residues). More structure-promoting amino acids could facilitate protein folding. Compared with disorder-promoting amino acids, all models designed sequences tended to have more disorder-promoting amino acids rather than CATH 100% (Fig. 2e).

### 2.3 The amino acids frequency of designed sequence

The frequency distributions of amino acid types for the designed sequences by different methods and for the native sequences were shown in Fig. 3a (14 *de novo* protein), Supplementary Fig. 5 (39 *de novo* proteins and single chains proteins). The Pearson correlation coefficient of the amino acid type compositions of the designed and the native sequences is 0.78, 0.80, and 0.81 on 14 *de novo* protein (Fig. 3b), 39 *de novo* proteins (Supplementary Fig. 5), and single chains proteins (Supplementary Fig. 5), respectively. The composition similarity of the amino acid type compositions of the designed and the native sequences is 0.42, 0.51, and 0.48. The ProteinMPNN reached the higher correlation (0.93) on 14 *de novo* protein, and the ESM-IF1 reached the higher correlation on 39 *de novo* protein (0.91) and on single chain protein (0.97).

**Fig. 3.**
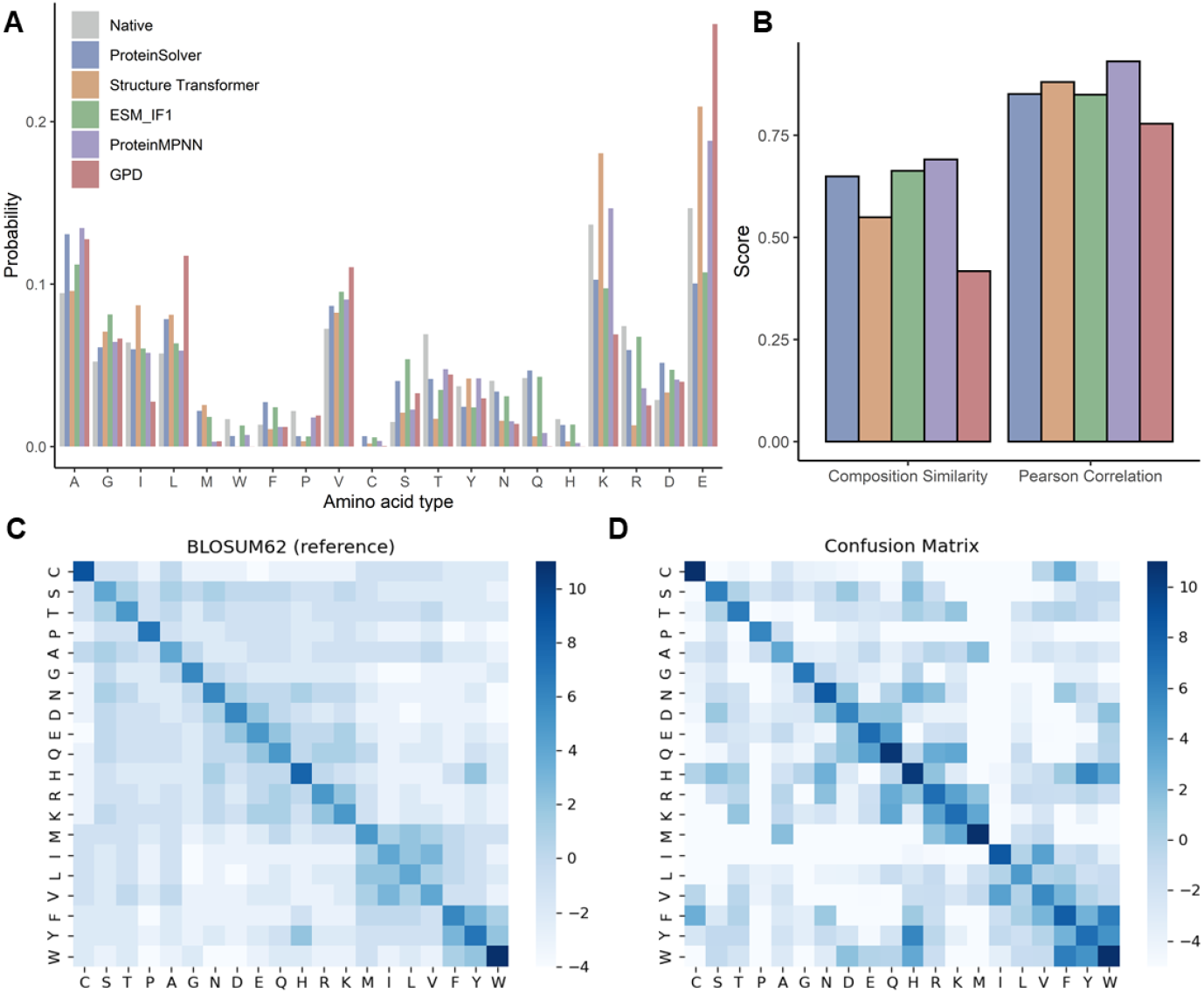
The amino acids frequency of designed sequence on 14 *de novo*. **a**, The sequence identity between the designed sequence and the native sequence of the target structure. **b,** The Pearson correlation coefficient and the composition similarity of the amino acid type compositions of the designed and the native sequences. **c,d,** Confusion matrix between native sequence and design sequences, compared to BLOSUM62 as reference.

Some sidechain types such as alanine, glutamicacid, leucine, valine have been used more frequently in the designed rather than the native sequences, while the sidechain types of obviously reduced usages in the designed sequences including isoleucine, threonine, asparagine, glutarnine, and arginine. The designed sequences has the higher frequency of non polar amino acids and the lower frequency of polar amino acids for all methods.

We calculated the substitution scores between native sequences and designed sequences by using the same log odds ratio formula as in the BLOSUM62 substition matrix (Fig. 3c and Fig. 3d). In the confusion matrixs, the diagonal elements have the biggest substitution scores for all amino acids, and the most amino acids of designed sequences physicochemically similar to the native types (for example, phenylalanine, tryptophan, and tyrosine; lysine, arginine, and methionine).

### 2.4 Experimental validation

Only a few studies so far have reported experimental examinations of protein sequences designed on the basis of deep learning. 3DCNN[15], ABACUS-R[16], and ProteinMPNN[18] designed sequences has got the experimentally solved atomic structures. ProteinSolver[13] and ProDESIGN-LE[21] designed sequences were shown to have desired secondary structure contents and to fold cooperatively according to circular dichroism signatures. None of these method analyzed the function and activity of designed proteins.

Here we examined the Candida antarctica lipase B (CalB) enzyme activity of designed sequences by GPD using wet experiment. CalB has been chosen to evaluate the GPD model because of its exceptionally robust tolerance to organic solvents and thermal stability, making it one the most commonly employed industrial enzymes for hydrolytic reactions in biocatalytic applications [25]. CalB belongs to α/β hydrolase family. The CalB has 317 amino acids and the total structure weight is 33.46 kDa. The CalB protein model was extracted from the Protein DataBank (pdb code: 1TCA). The substrate is p-Nitrophenyl acetate C2.

#### 2.4.1 CalB design

CalB has the catalytic triad formed by residues S105, D187 and H224. The activesite cavity is shaped as a tunnel that limits the steric positioning of substrates. 62 sites residue positions including 5 active-sites amino acids, 19 substrates pocket amino acids, 20 conserved sites from CalB single site-saturation mutagenes data, and 18 conserved sites from multiple sequence alignment, were fixed (see method). We used GPD to generate 1,000,000 *de novo* designed sequences of CalB.

#### 2.4.2 Functional screening

The functional screening of 1,000,000 CalB designed sequences was shown in Fig. 4a. The designed sequences were virtually screened on the basis of protein folding ability, protein solubility, and molecular dynamics (MD) simulation.

**Fig. 4.**
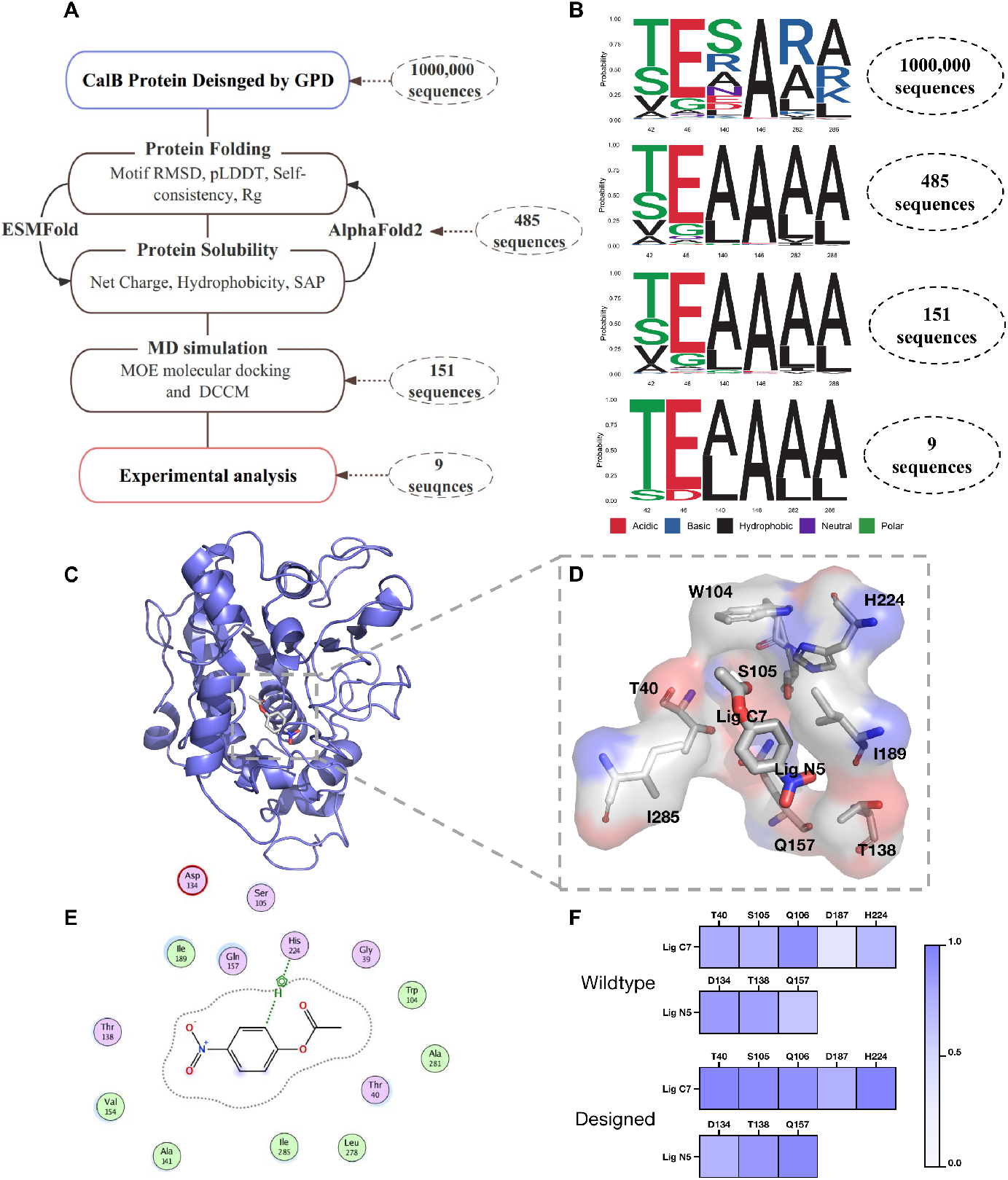
The design workflow of CalB hydrolase. **a**, The design workflow of CalB including CalB sequences design, protein folding ability, protein solubility, MD simulation. **b,** The seqlog plot of Thr42, Gln46, Leu140, Ala146, Ala282, Val286 after each step screening. **C,** The CalB and substrate (p-Nitrophenyl acetate C2) complex after MD simulation. **d,** The region of CalB active sites is enlarged. **e,** The 2D interaction diagram between CalB and substrate. **f,** The Dynamic Cross-Correlation Matrix (DCCM) characterize the significant interactions of CalB active sites and substrate.

**Protein folding ability**. We applied ESMFold and AlphaFold2 to predict the structures of the 1,000,000 designed sequences. The RMSD of 62 conserved site between the predicted structures and CalB native structures, the pLDDT scores, self-consistent between ESMFold and AlphaFold2, and the radius of gyration (Rg) between the predicted structures and CalB native structures were used to estimate the folding ability of designed sequences.

**Protein solubility**. The net charge, hydrophobicity, and Spatial Aggregation Propensity score (SAP) was used to estimate the protein solubility. 151 designed sequences were selected after protein folding and solubility screening.

**MD simulations**. MD simulations were carried out for the 151 protein-ligand complexes from MOE docking result. 9 sequences meet the catalytic mechanism and were selected for experimental validation. The RMSD of 9 sequences ranges from 2.29-3.38^°^A and the recovery ranges from 0.445-0.498. The diversity between 9 sequences were 0.215-0.253.

The complete data of each step screening were reported in supplemantary Table. 1. Fig. 4b showed the CalB enzyme active sites. Fig. 4c and Supplementary Fig. 6 show the seqlog plot of import residues after each step screening, indicating that our virtual screening workflow is able to select the residues that meet the chemistry properties. Thr42, Gln46, Leu140, Ala146, Ala282, Val286 were near by the conserved sites.

#### 2.4.3 Experimental validation

9 sequences meet the catalytic mechanism and were selected for experimental validation(Supplementary Fig. 7). Among these 9 designed sequences, 5 have led to successful protein expression in Yeast. All of the expressed proteins could be purified in soluble form. Two designed sequences (CalB D263 and CalB D323) have the activity (Fig. 5a and Fig. 5b). The high success rate of the experiments indicates the efficiency of our design and screening process.

**Fig. 5.**
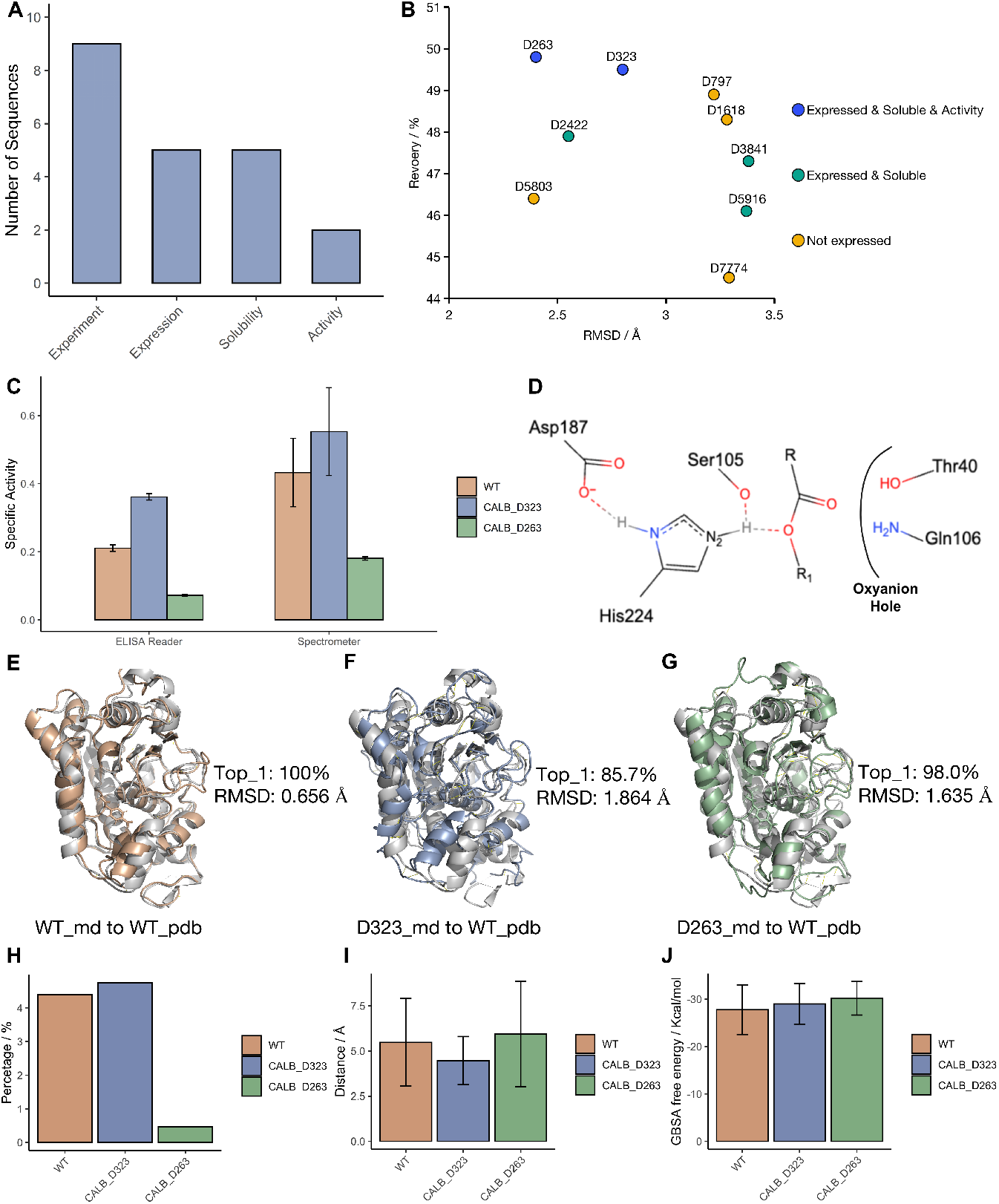
Experiments and MD simulation results of designed sequences. **a**, The number of designed sequences are experimented, and expressed. **b,** The RMSD and recovery values of designed sequences. **c,** The specific activity of designed sequences and wild type using Spectrometer and ELISA Reader. **d,** The model of pre-catalytic state of CALB-substrate complex. **e,** The alignment between MD simulation clustered WT structure and WT pdb structure. **f,** The alignment between MD simulation clustered D323 structure and WT pdb structure. **g,** The alignment between MD simulation clustered D263 structure and WT pdb structure. **h,** The hydrogen bond percentage during the MD simulation trajectories. **i,** The distance between Ser105 side-chain gamma oxygen atom and the carbonyl carboon atom. **j,** The GBSA binding free energy.

As Fig. 5b shows, the two catalytical positive proteins have lower RMSD and higher recovery, compared to the negative ones. This result shows that our screening work-flow could somehow be useful.

The purified proteins have been subjected to specific activity analysis using Spectrometer and ELISA Reader. The specify activity measured by ELISA Reader was 0.210±0.0065 U/mg, 0.361±0.0089, and 0.072±0.0029 for CalB native sequence, CalB D323, and CalB D263, respectively. The designed sequences (CalB D323) has the higher hydrolytic activity than CalB native sequence (P value is 0.029). The specify activity measured by Spectrometer was 0.433±0.1005 U/mg, 0.553±0.1288, and 0.1807±0.0053 for CalB native sequence, CalB D323, and CalB D263, respectively. The hydrolytic activity between the CalB D323 and CalB native sequence (P value is 0.029) has no significant difference (P value is 0.1). The wet experiment demostrated that the CalB *de nono*designed sequnce has the higher activity than CalB native sequence.

To further understand the experimental result, we performed the MD simulation for wild type, D323 and D263, respectively. For each system, we ran three parallel trajectories of 200ns. Fig. 5e to g shows that the designed proteins have higher RMSD to wild type crystal structure. However, the RMSD of simulated structure is much lower than the AlphaFold2 predicted one (See in Fig. 5b), indicating that MD simulation may be a way to enhance the structure prediction result. As Fig. 5d shows, the hydrogen bond between Ser105 and His224 is essential to CALB’s catalytic ability. Fig. 5h shows that the wild type and D323 protein have much higher percentage of hydrgen bond compared to D263. This may be one of the explanation of the experimental result. Meanwhile, the distance between Ser105 side-chain gamma oxygen atom and the ligand carbonyl carboon atom of D323 is lower than that of wild type and D263, shown in Fig. 5i This may also shows why D323 has the highest catalytic ability among the three. However, the GBSA results in Fig. 5j shows no significant difference among three systems. Hence, the GBSA results may only show the binding affinity, but not the catalytic capability.

### 2.5 GPDGenerator Webserver

As shown in Supplementary Fig. 8, we also developed a user-friendly online tool for the GPDGenerator (https://yu.life.sjtu.edu.cn/ChenLab/GPDGenerator/) and made it publicly accessible. The server uses the parameters derived from the trained model to design protein sequences from the input PDB file.

The “Introduction” interface introduce what is GPD and shows examples of GPD results. At the “Analysis” interface, the users input the PDB file and could design the amino acids of interest on specified positions. The “Result” interface output the designed sequences as the FASTA format, and output the recovery with the corresponding native sequence. The protein length should be less than 400 amino acids. The number of designed sequences should be less than 100 for saving computer resources. In summary, this online tool should facilitate the generation of novel protein sequences based on fixed-backbone.

## 3 Discussion

This study devised a graph representation-based Transformer to address the fixed-backbone protein sequence design. Compared with existed deep learning protein sequence design approaches, the main contributions of our GPD model are two aspects. Firstly, the GPD model incorporated five graph node encodings and four edge encodings, and is able to leverage the protein spatial information. We added a normally distributed random matrix in node features to increase the diversity of designed sequences. Secondly, functional filtering based on the structure folding, solubility, and function were performed to improve the success rate in experiments. GPD designed sequences of CalB hydrolase has the higher specify activity than CalB wild type.

The time consuming of GPD is acceptable. The time consuming for designing 100 sequences with 261 residues by CPU were 3 minutes, 25 seconds, 33 minutes, 112 seconds, 6.25 days, 2.86 days, and 35 seconds for ProteinSolver, Structure Transformer, ESM-IF1, ProteinMPNN, 3D CNN, ABACUS-R, and GPD, respectively. Compared with three methods (3DCNN, ABACUS-R, and ProteinMPNN) with experimental examinations by crystallograph, our GPD model spend less time consuming. Our GPD model is more suitable for high throughput sequence generation.

GPD achieve higher recovery and diversity on different type proteins. Existed methods only sought to improve the protein sequence recovery, however, the native recovery rate may not be pursued as a “golden standards” for benchmark of different methods. Proteins consist of any combination of 20 amino acid residues, so the sequence space is vast. Ideally, the designed sequences should cover a wide range of sequence space landscape and have higher diversity.

High recovery are still insufficient for predicting the performance of design methods in wet experiments. The high recovery have not indicated substantial correlations with the success rate of wet experiment (a single residue substitution that does not cause any notable changes in such metrics can be enough to disrupt the overall structure). Designed proteins could be expressed and purified is import for the wet experiment. Functional filtering based on the structure folding, solubility, and function are essential for improving the success rate of wet experiment. Structure folding and solubility constitutes useful protocols for computationally assessing the design sequences.

We used GPD to design 1,000,000 de novo sequences of CalB hydrolase. Nine sequences after functional filtering were examined by wet experiments. More than half (5/9) of the designed sequences could be expressed and purified. Two of nine experimentally examined designed sequences of CalB has hydrolytic activity, and The specify activity of one designed sequences (0.36) was significant higher than CalB wild type(0.21). The high success rates of experimental design of GPD, together with the compute efficiency and lack of requirement for customization, make GPD very broadly useful for protein design.

Although our method has achieved a satisfactory performance on fixed-backbone protein sequence design, there are also several ways for further improving. Firstly, the protein folding ability of designed sequence is limited (Table 1). The highest percentage of designed sequences with RMSD less than 2^°^A is only 23.6% for single chain proteins, and all methods still have the deficiencies of protein folding. Secondly, the number of expressed and soluble proteins are not very high compared with ProteinMPNN and ABACUS-R. The ratio of expressed and soluble protein are 76% and 86% for Protein-MPNN and ABACUS-R, respectively. The ratio is only have 56% for GPD. Thirdly, the fixed-backbone sequence design for different type proteins should be validated by wet experiments in the future studies.

**Table 1.**
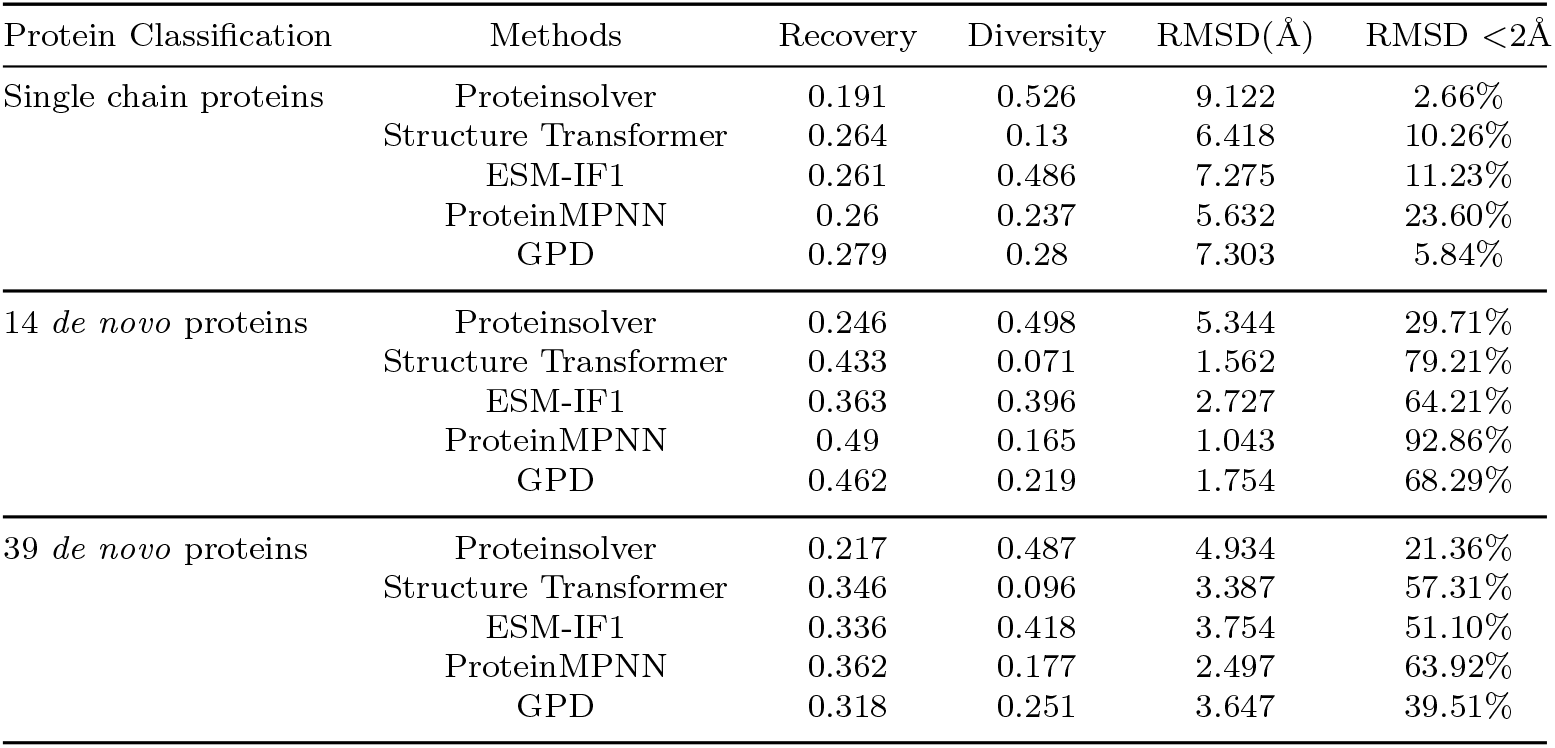
The performance of different methods.

| Protein Classification | Methods | Recovery | Diversity | RMSD(Å) | RMSD < 2Å |
| --- | --- | --- | --- | --- | --- |
| Single chain proteins | Proteinsolver | 0.191 | 0.526 | 9.122 | 2.66% |
|  | Structure Transformer | 0.264 | 0.13 | 6.418 | 10.26% |
|  | ESM-IF1 | 0.261 | 0.486 | 7.275 | 11.23% |
|  | ProteinMPNN | 0.26 | 0.237 | 5.632 | 23.60% |
|  | GPD | 0.279 | 0.28 | 7.303 | 5.84% |
| 14 *de novo* proteins | Proteinsolver | 0.246 | 0.498 | 5.344 | 29.71% |
|  | Structure Transformer | 0.433 | 0.071 | 1.562 | 79.21% |
|  | ESM-IF1 | 0.363 | 0.396 | 2.727 | 64.21% |
|  | ProteinMPNN | 0.49 | 0.165 | 1.043 | 92.86% |
|  | GPD | 0.462 | 0.219 | 1.754 | 68.29% |
| 39 *de novo* proteins | Proteinsolver | 0.217 | 0.487 | 4.934 | 21.36% |
|  | Structure Transformer | 0.346 | 0.096 | 3.387 | 57.31% |
|  | ESM-IF1 | 0.336 | 0.418 | 3.754 | 51.10% |
|  | ProteinMPNN | 0.362 | 0.177 | 2.497 | 63.92% |
|  | GPD | 0.318 | 0.251 | 3.647 | 39.51% |

The ProteinMPNN designed sequences have the higher protein folding ability than other methods, even so, the percentage of designed sequences with RMSD less than 2^°^A is only 23.6% for single chain proteins. All these methods still have the deficiencies of protein folding, and this will lead to experimental failure. The protein folding ability is the important metrics to evaluate the performance of different methods.

## 4 Methods and Materials

### 4.1 Feature representation

In order to obtain as much structure information as possible and satisfied the SE(3) equivariance, we treated each single protein main-chain structure as a graph that contains both node features and edge features. We took every single residue as a node and took the connection between residues as the edge of the graph. All node features and edge features could be computated by only the backbone atom information and satisfied the SE(3) equivariance.

The node features contained the main-chain dihedral angles psi and phi, the secondary classification, the centrality of each residue, the pre-designed protein sequence, and a tensor of random seed. For the main-chain dihedral angle, both phi and psi were embedded in the ways of sine and cosine function (shown in Figure 1A). This information redundancy could help the neural network better learned the features. We used the Define Secondary Structure of Proteins (DSSP) algorithm to classify the main-chain secondary structure. Eight class of secondary structures were used in this study, such as 310-helix, α-helix, π-helix, hydrogen bonded turn, β-harpin, β-bridge, bend, and loop. The centrality of a single residue was represented by the betweenness centrality and calculated by

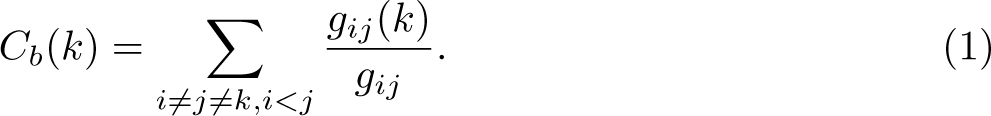

where c*_b_*(k) means the betweenness centrality of node k. g*_ij_* means the number of shortest paths that start from node i then end with node j, and at the same time pass node v*_i_*. g*_ij_* means the number of all shortest paths between node i and j.

For the network was trained for protein sequence design, we leaved an API for predesigned sequence embedding. The user could pre-design each residue at each position and the other residues will be generated according to both the structure information and the pre-designed sequence. If there is no pre-designed residue requirement, the pre-designed sequence tensor would be set as all-zeros. Taken Figure 1B for example, when redesigning GFP, we could predefine the chromophoric residues that could not be predicted by the backbone structure but necessary for GFP function (citation for GFP).

The random tensor was used for enlarging the designed sequence space, which is of essential importance of de novo protein design. Our goal is to get a higher level of neural network randomicity when training at the same level of loss value.

The edge features contained the distances, the movement vectors, the shortest pathway, and the rotation quaternions. For the distances map, we calculated the distances between the alpha carbon atoms of each residue (shown in Figure 1D). For the movement vector, we used the coordinate system transformation to satisfy the SE(3) equivariance. First, by using the residue gas shown in Figure 1D, (citation for residue gas) we defined a residual specific coordinate system O, defined as:

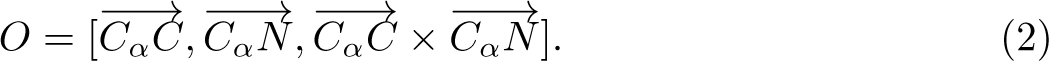

The residual specific coordinate system based on movement vector v*_m_* is calculated by:

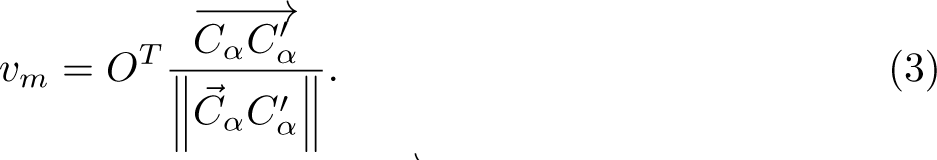

where *v_m_* means the transferred movement vector, 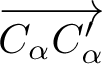 means the initial movement vector under the Cartesian coordinate system.

For the rotation relation between two residues, we used rotation quaternion for representation, defined as:

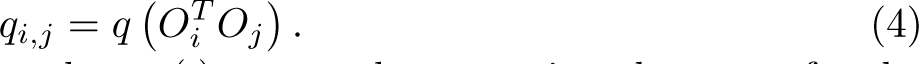

where q*_i,j_*means the quaternion number, q(·) means the operation that transfer the rotation matrix into rotation quaternion number. The O*_i_* and O*_j_* are the two residual coordinate system representation of residue i and j, respectively.

The shortest pathway feature was calculated based on the distance matrix, also a way of feature redundancy. We set the cut of 12^°^A between two carbon alpha atoms to define whether there is a contact between two residues. By using this contact map, we calculates the length of the shortest path between each residue pairs, see Figure 1E for example.

### 4.2 Model structure

The overall structure of GPD was shown in Figure 2A, using only the encoder part of the real Graphormer for save more computational resources. Six recycles of the Graphormer attention block were used. The designed sequences were brought out by a linear layer continue with the softmax operation.

The key component of Graphormer was the Graphomer block and shown in Figure 2B. The main difference between Graphormer and traditional Transformer was that the adding of the embedded edge feature. In every Graphormer block, the interaction weight between two residues’ node features was determined by both the attention matrix and the edge feature matrix. This allowed the structural information flows from edges to nodes. Also, the node information could flow to edges in the second Graphormer block.

The embedding blocks for edge features and node features were shown in Figure 2C and Figure 2D, respectively. The Distances map, the movement vector matrix, the rotate quaternion matrix and the shortest path number matrix were all join at the last dimension and passed through two layer of fully connected neural networks. The last dimension of the edge features matrix was transferred into the exact dimension of the number of the attention heads in the multi-head attension block.

### 4.3 Data set and benchmark metrics

We used the CATH 40% sequential non-redundancy dataset for neural network training, validation and testing. The split radio between training, validating and testing set was 29868:1000:103. We trained the network for 400 epochs with the randomly masked pre-designed sequence, and validated with the fully masked pre-designed sequence. 14 de novo protein, 39 de novo proteins, and 103 single chains proteins were used to evaluate the performance of GPD model and other existed methods.

Recovery was the proportion of the same amino acids at equivalent position between the native sequence and the designed sequence, calculated with Eq. 5. Diversity was one minus the proportion of the same or similar amino acids at equivalent position between designed sequence, calculated with Eq. 6. RMSD quantifies the differences between the predicted structures and the corresponding native structures. RMSD is calculated as Eq. 7.

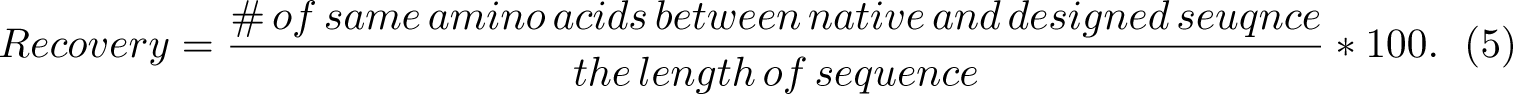

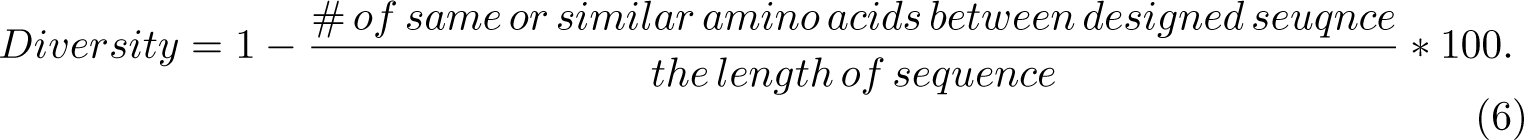

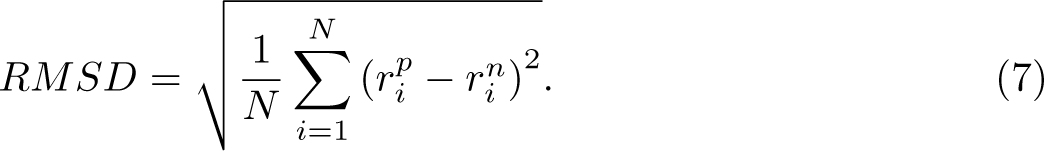

where *r^p^_i_* and *r^n^_i_* are Cartesian coordinates of the i-th atom from predicted structure *r^p^* and the native strcture structure r*^n^*, respectively. N is the number of atoms.

### 4.4 CalB design

62 sites residue positions were fixed according to the enzyme catalytic mechanism. Residues S105, D187, H224 were catalytic triad; T40 and Q106 were oxyanion hole; A141, L144, V149, and I285 were subtrate hydrophobic pocket; D134, T138, and Q157 were subtrate hydrophilic pocket. 20 conserved residues (38, 39, 107, 108, 109, 110, 111, 133, 180, 181, 182, 190, 209, 230, 79, 130, 131, 132, 135, 228) were from CalB single site-saturation mutagenes data. 18 conserved residues (103, 104, 190, 69, 180, 209, 216, 239, 188, 258, 150, 136, 294, 127, 74, 98, 169, 64) were from multiple sequence alignment. MD simulation showed that 15 residues (268-280, 225, 154) were important to limit the steric positioning of substrates. These 62 conserved residues of CalB were fixed when we used GPD to design 1,000,000 *s*equences .

### 4.5 Functional screening

The designed sequences were virtually screened on the basis of protein folding ability, protein solubility, and molecular dynamics (MD) simulation. According to the structure of CalB, 7 residues (277, 280, 281, 285, 139, 188, 38) near by subtrate hydrophobic pocket should be non-polar. 40278 designed sequenced were non-polar amino acid on these 7 residues. ESMFold was used to predict the structure of these sequences. Firstly, We applied ESMFold to predict the structures of the 40278 designed sequences. 485 sequences meet the meet the screening criteria of protein folding ability and protein solubility. Secondly, we used AlphaFold2 to predict the structures of these 485 sequences, 151 sequences meet the meet the screening criteria of protein folding ability and protein solubility. MD simulations were carried out for the 151 protein-ligand complexes. 9 sequences meet the catalytic mechanism and were selected for experimental validation.

#### 4.5.1 Protein folding ability

The radius of gyration (Rg) of C*_α_*, the RMSD of 62 conserved site (Eq. 7), and the pLDDT scores were used to estimate the folding ability of designed sequences. Rg of C*_α_* determines the compactness of predicted structures, smaller means that the protein structure is more compactness and stable. The Rg is calculated by mdtraj [26, 27], and the Rg of designed proteins should be less than CalB native structures (18.45^°^A). RMSD the quantifies the differences of 62 conserved site between the predicted structures and CalB native structures. The RMSD of 62 conserved site should less than 1.5^°^A and the pLDDT of predicted structures should great than 80. The Rg only measures the folding ability of predicted proteins as a whole, while the the RMSD of 62 conserved site guarantees the similarity of activity sites.

#### 4.5.2 Protein solubility

The net charge, hydrophobicity, and SAP was used to estimate the protein solubility. The net charge of a protein is important for its solubility, and neutral or positively charged proteins are more likely to lead to aggregation, and neutral or positively charged proteins may have non-specific binding with negatively charged DNA (Eq. 8). SAP was calculated for each residue by a combination of solvent accessibility area and hydrophobicity, calculated by Rosetta. Hydrophobicity control the non-polar residues on the surface (Eq. 9, 10) [28].

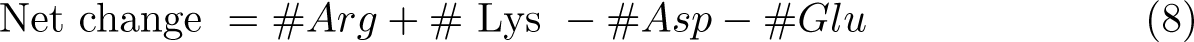

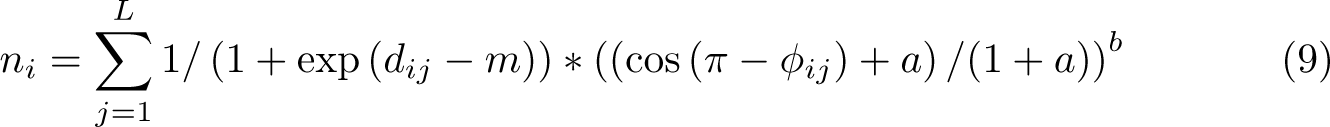

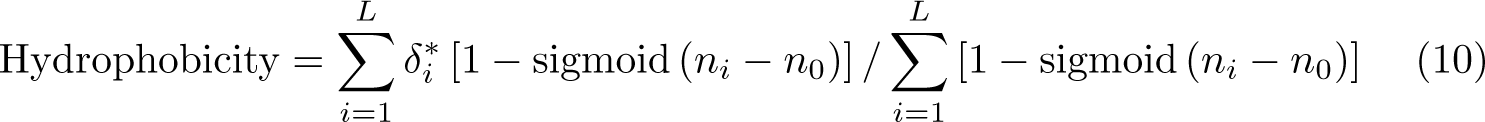

where d*_ij_* and ϕ*_ij_* are the C*_b_* − C*_b_* distance and Ca − Cb/Ca − Cb angles between residues i and j, and m=1, a=0.5, and b=2 are tuning parameters set to their default values. n_0_ was the median of n*_i_*. δ*^∗^_i_* = 1 if residue i is non-polar (V, I, L, M, W, F) and 0 otherwise. The quantity 1 − sigmoid (n*_i_* − n_0_) ranges from 0 to 1 and is higher when a residue is closer to the surface. More nonpolar residues on the surface would disrupt protein folding.

#### 4.5.3 MD simulations

MD simulations were carried out for the 151 protein-ligand complexes from MOE docking result. Firstly, energy minimization, heating and equilibrium of the system are carried out. The energy of the system is minimized by the steepest descent method of 3000 steps and the conjugate gradient method of 3000 steps. After energy minimization, the system is heated from 0K to 321K in a time of 50ps, and then performs an energy balance of 100ps at constant pressure and temperature of 321K. In the whole process, the long range electrostatic interaction is calculated by PME algorithm, and the covalent bonds of all hydrogen atoms are constrained by SHAKE algorithm. The cut-off value for the van der Waals interaction and the short-range electrostatic interaction is set at 8^°^A. The final simulation process was carried out at NPT and temperature of 321K, and the simulation time was 20ns. 9 sequences with reasonable conformation were selected for experimental validation. After experiments, MD simulation of the wild type complex, the D323 and the D263 were performed with three parallel trajectories for 200ns.

#### 4.5.4 Trajectory analysis

The Dynamic Cross-Correlation Matrix were calculated as follows:(Eq. 11)

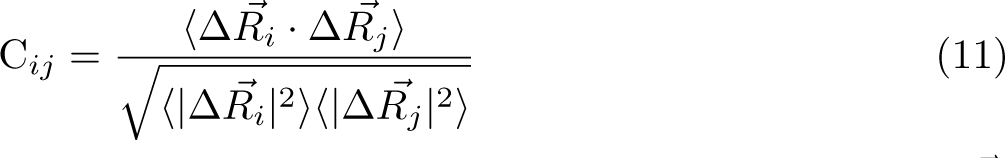

where C*_ij_* is the cross-correlation of atom i and atom j, ⟨⟩ denotes time averaging, Δ^R⃗^*_i_* and Δ^R⃗^*_j_* represent the displacement of atom i and j respectively. When calculating DCCM, the python package MDtraj was used for trajectory loading [27]. The cpptraj was used for RMSD, distance, hydrogen bond and structure cluster analysis. Clustering uses the DBSCAN method [29], taking 1 frame every 100 frames. MinPoints is set to 10, and epsilon is set to 3.0. When calculating the hydrogen bond, the distance cutoff was set to 0.3nm, while the angle cut-off was set to 120 degree. The binding free energies of the protein-ligand complexes were computed using the Generalized Born Surface Area (GBSA) model [30].

### 4.6 Heterologous expression and purification of CalB

The Escherichia coli Rosetta (DE3) with the recombinant plasmid of pET22b-CalB designed sequences was cultivated for 3h at 37 ℃ in 2×yeast extract tryptone (2YT) with ampicillin (100 µg/mL) and chloramphenicol (34 µg/mL). Then the final concentration of 0.1 mM IPTG (Isopropyl-D-Thiogalactoside) was added and induced overnight at 15℃. Cells were collected by centrifugation at 5000 rpm for 10 min. The designed proteins were purified by Nickel column affinity chromatography. The purified proteins ware detected by sodium dodecyl sulfate polyacrylamide gel electrophoresis (SDS-PAGE). The recombinant plasmid of pET22b-CalB was synthesed in GENEWIZ Company (Suzhou, Chian).

### 4.7 Enzyme activity assays

The p-Nitrophenyl acetate C2 was used to determine the activity of CalB designed sequences and CalB wild type. The ability of enzymatic hydrolysis was measured by UV-2550 ultraviolet spectrophotometer (Shimadzu, Kyoto, Japan) at 405 nm wavelength and ELISA. The reaction system consisted of p-Nitrophenyl acetate C2 (200 µM) and 100 µL enzyme solution, and the reaction mixture was supplemented to 1 mL by 50mM PBS (pH 7.5). The enzymatic reaction was carried out at 37°C for 5 minutes. One unit of enzymatic activity (U) was defined as the amount of enzyme required to hydrolyze the substrate to produce 1 mol of p-nitrophenol per min. p-Nitrophenyl acetate C2 (CAS No. 830-03-5) was from Sigma-Aldrich (St.Louis, MO, USA).

## Appendix A Supplementary Information

**Fig. A1.**
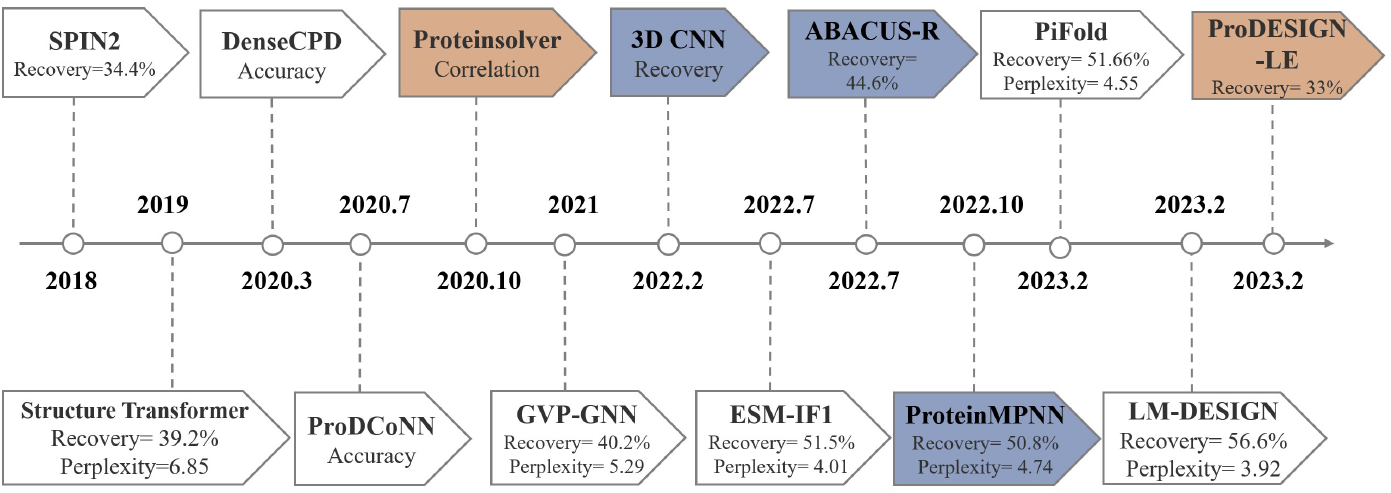
The deep learning-based protein sequence design methods.

**Table A1.**
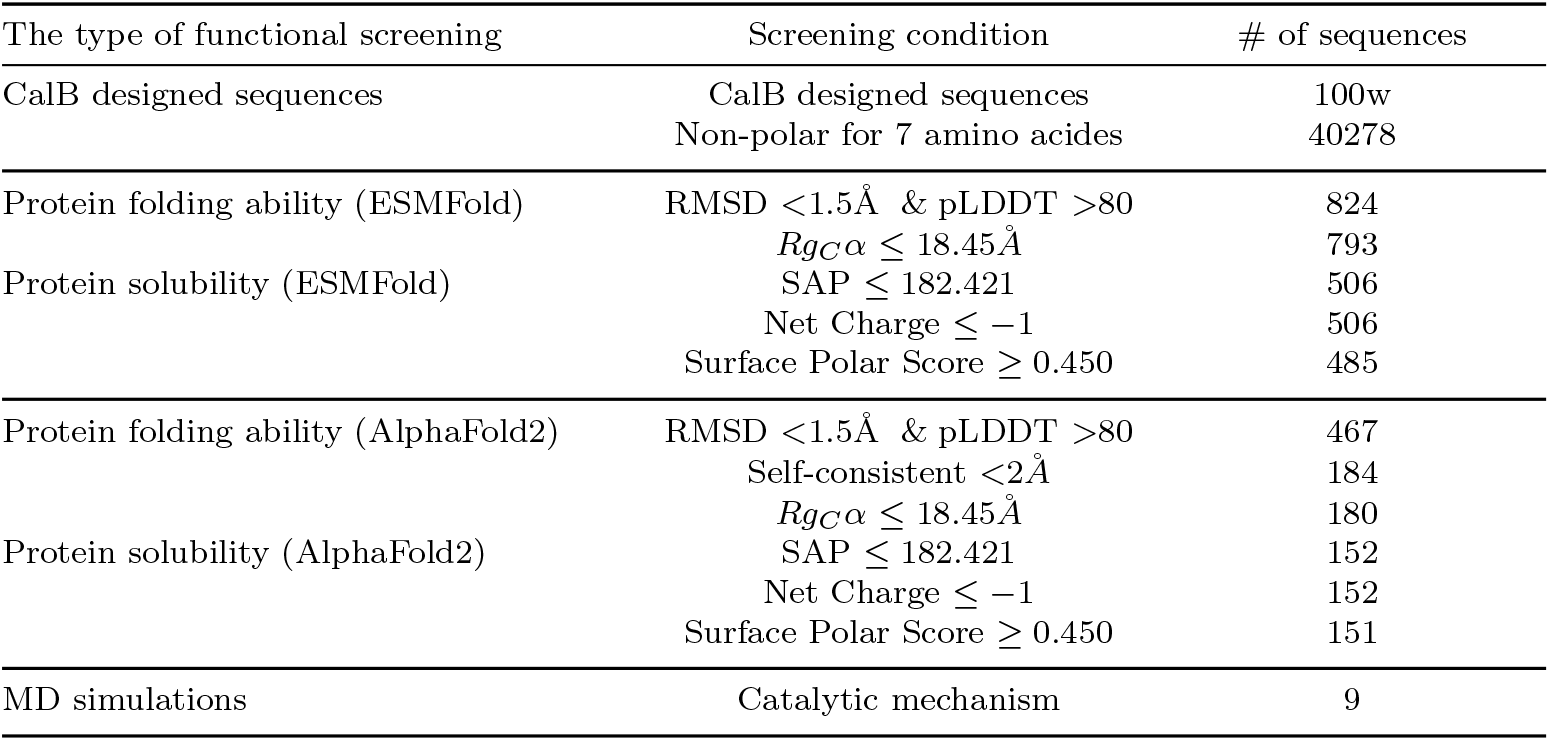
The functional screening.

**Fig. A2.**
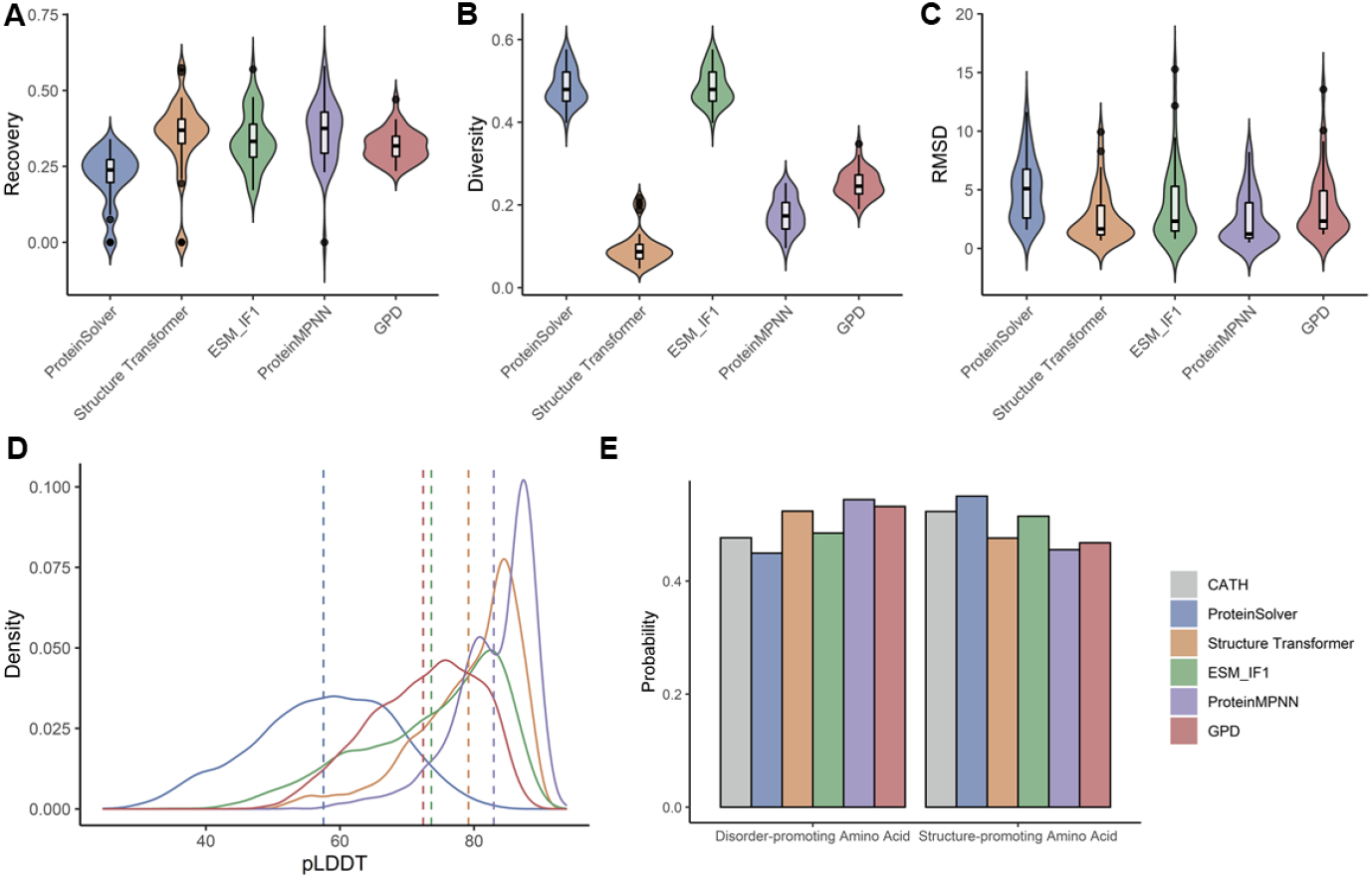
Comparison of designed sequence for five methods on 39 *de novo* proteins. **a**, the sequence recovery between the designed sequence and the native sequence of the target structure. **b,** The diversity of designed sequences. **c,** RMSD for aligning the ESMFold predicted structures with the corresponding native structures. **d,** The pLDDT scores of the ESMFold predicted structures. **e,** the frequency of disorder-promoting amino acids (alanine, glycine, proline, arginine, glutamine, serine, glutamic acid, and lysine) and structure-promoting amino acids (other twelve residues).

**Fig. A3.**
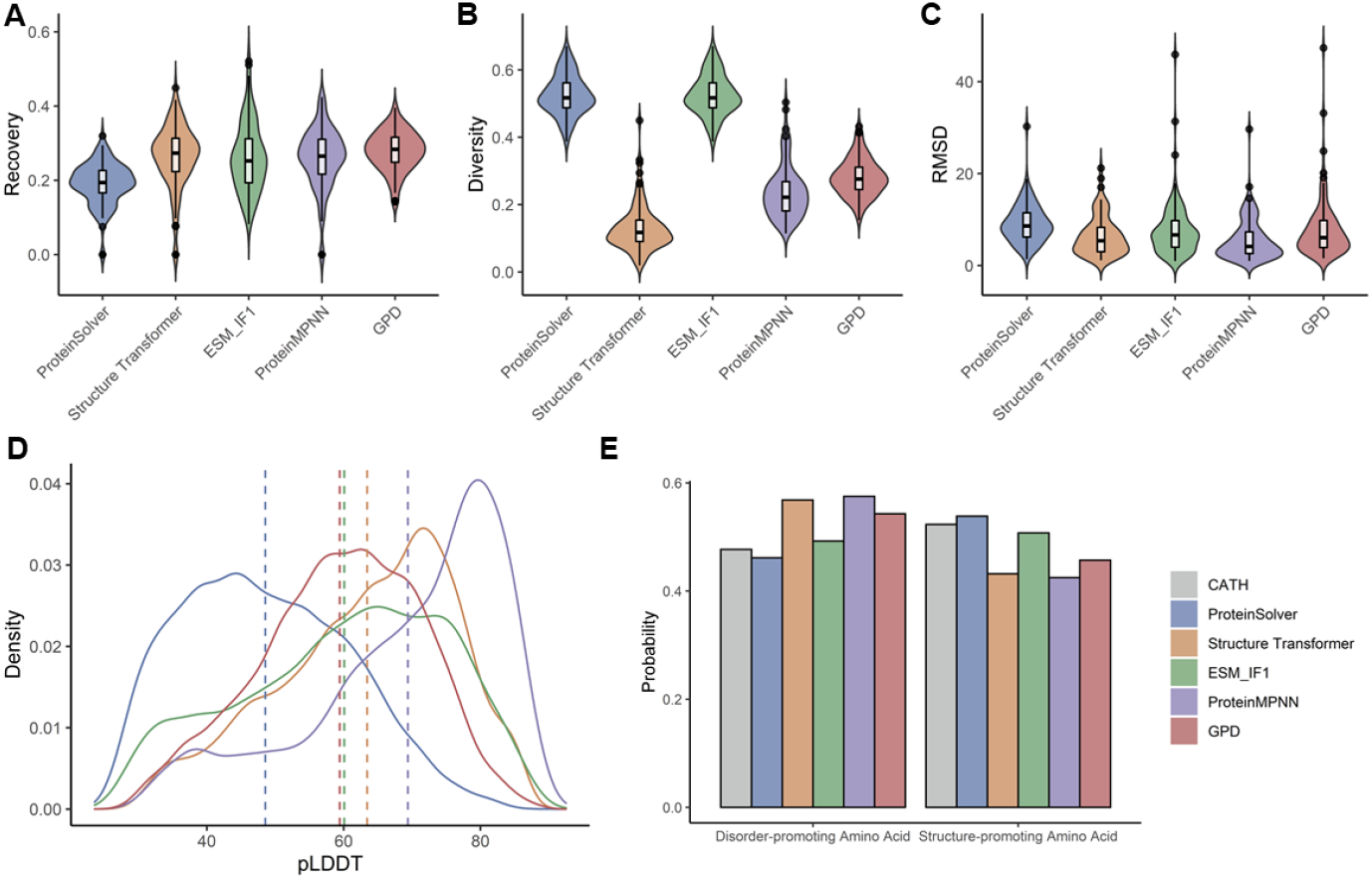
Comparison of designed sequence for five methods on 103 single chain proteins. **a**, the sequence recovery between the designed sequence and the native sequence of the target structure. **b,** The diversity of designed sequences. **c,** RMSD for aligning the ESMFold predicted structures with the corresponding native structures. **d,** The pLDDT scores of the ESMFold predicted structures. **e,** the frequency of disorder-promoting amino acids (alanine, glycine, proline, arginine, glutamine, serine, glutamic acid, and lysine) and structure-promoting amino acids (other twelve residues).

**Fig. A4.**
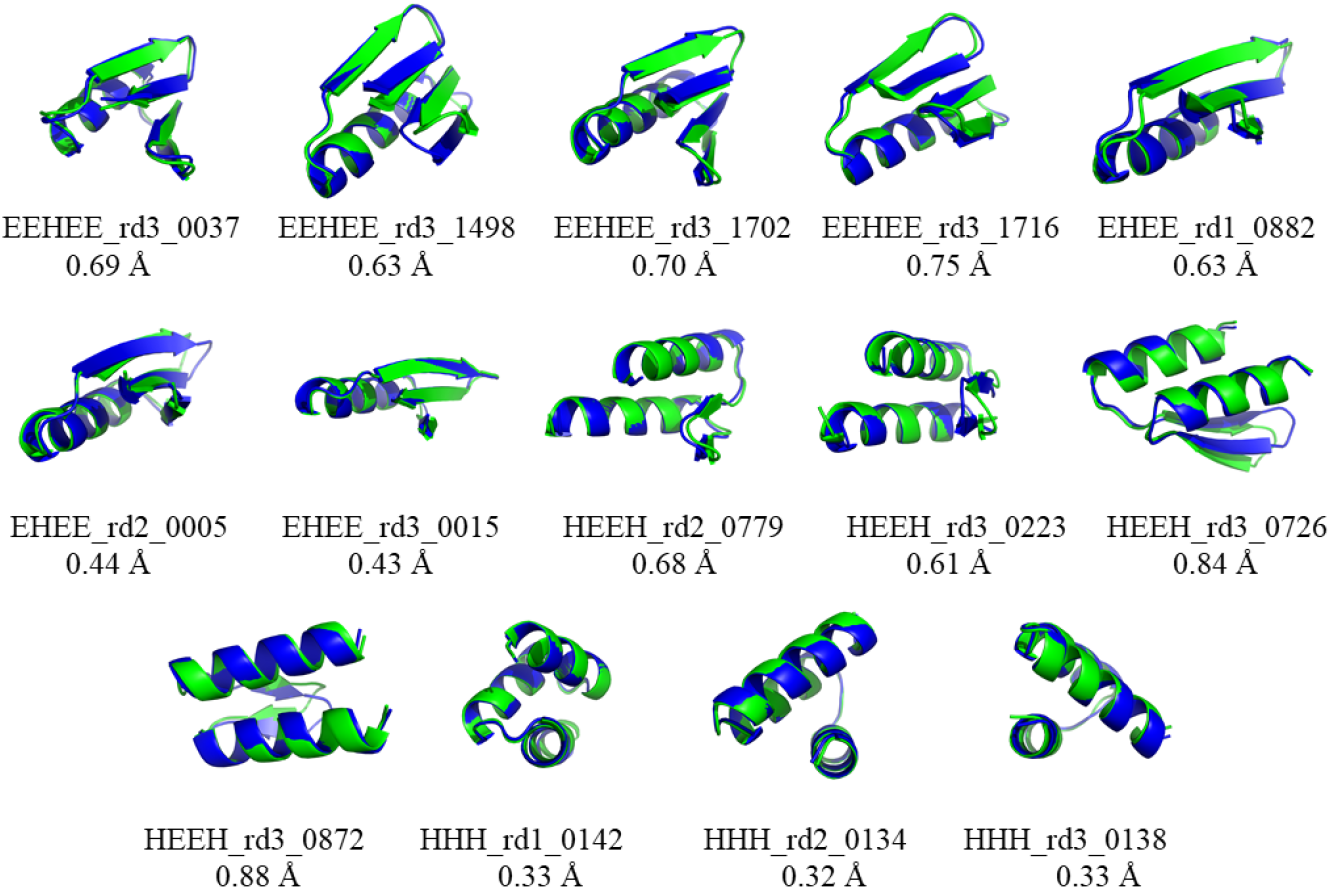
The amino acids frequency of designed sequence on 39 *de novo* (a,b) and on 103 single chian proteins (c,d). **a, c**, The sequence identity between the designed sequence and the native sequence of the target structure. **b, d,** The Pearson correlation coefficient and the composition similarity of the amino acid type compositions of the designed and the native sequences.

**Fig. A5.**
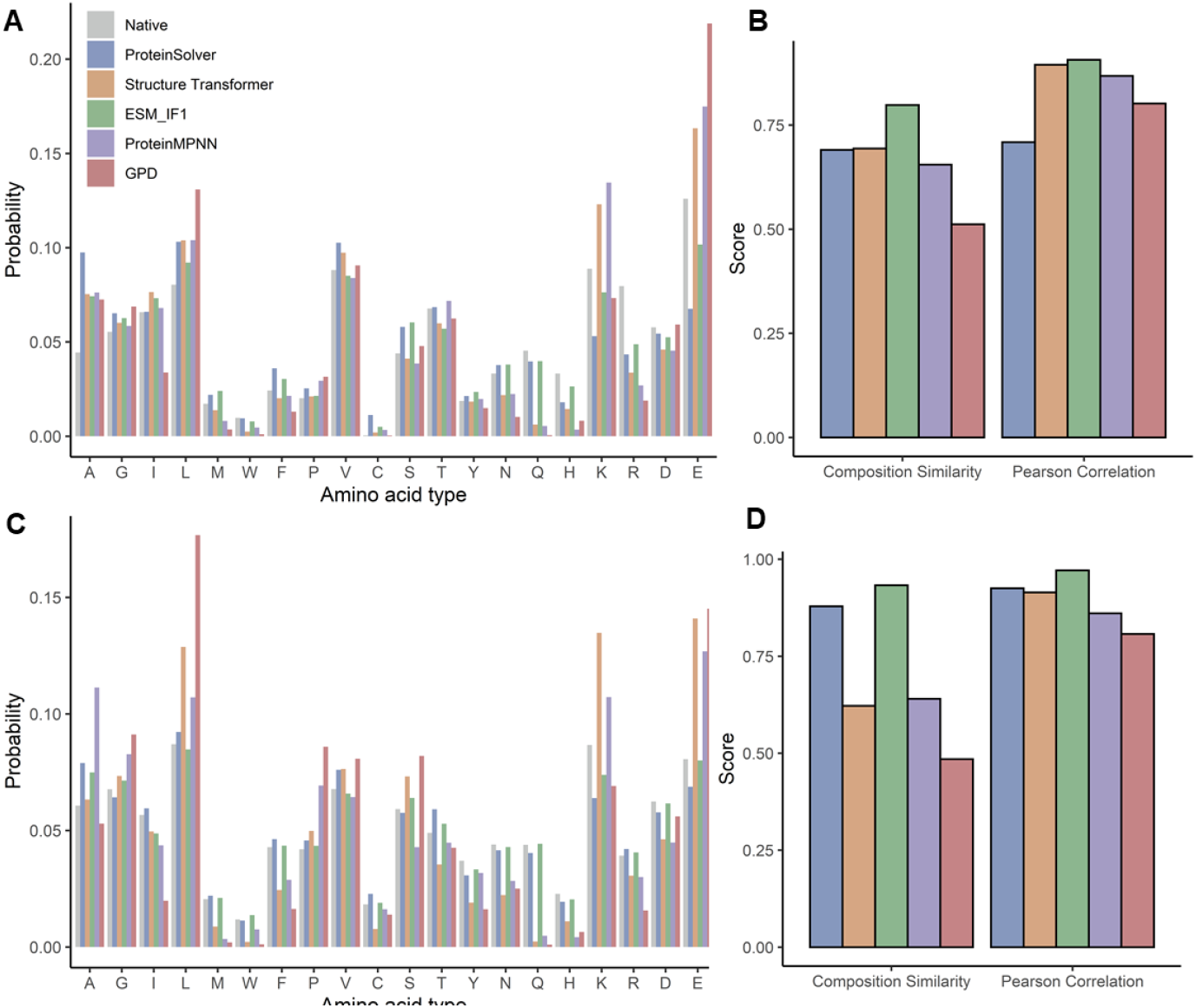
Structure prediction by AlphaFold2 on designed sequences. Overlay of native structures (green) with AlphaFold2 predicted structures of GPD model designed sequences (blue). The predicted structure with the minimum RMSD was shown.

**Fig. A6.**
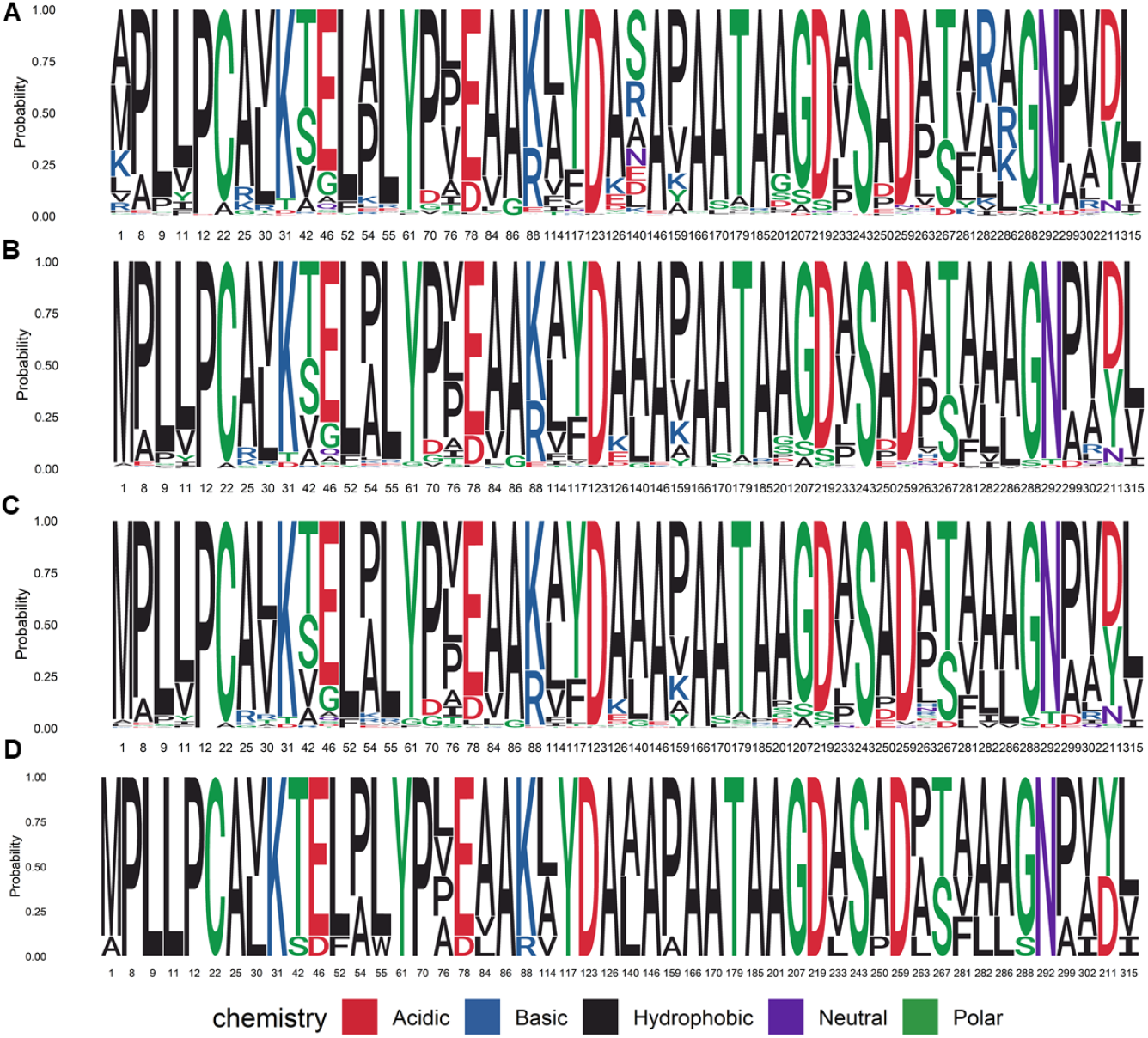
The seqlog plot after each step screening. **a**, The seqplot of 50 residues of 1,000,000 designed sequences of CalB. **b,** 485 sequences meet the meet the screening criteria of protein folding ability and protein solubility based on ESMFold. **c,** meet the meet the screening criteria of protein folding ability and protein solubility based AlphaFold2. **d,** The seqlog plot of 9 sequences meet the catalytic mechanism and were selected for experimental validation.

**Fig. A7.**
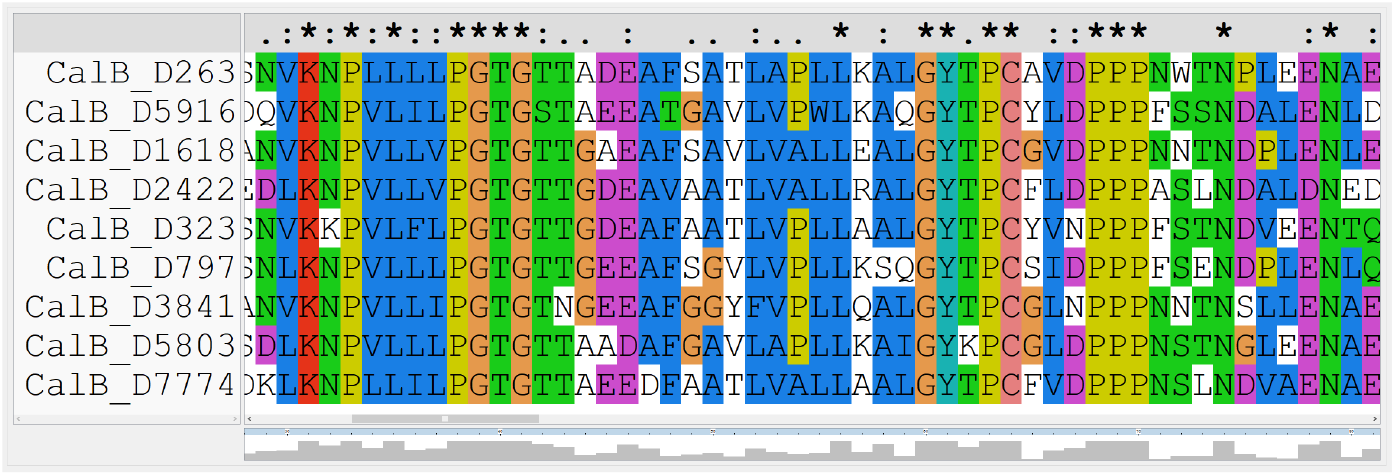
The 9 sequences for experimental validation.

**Fig. A8.**
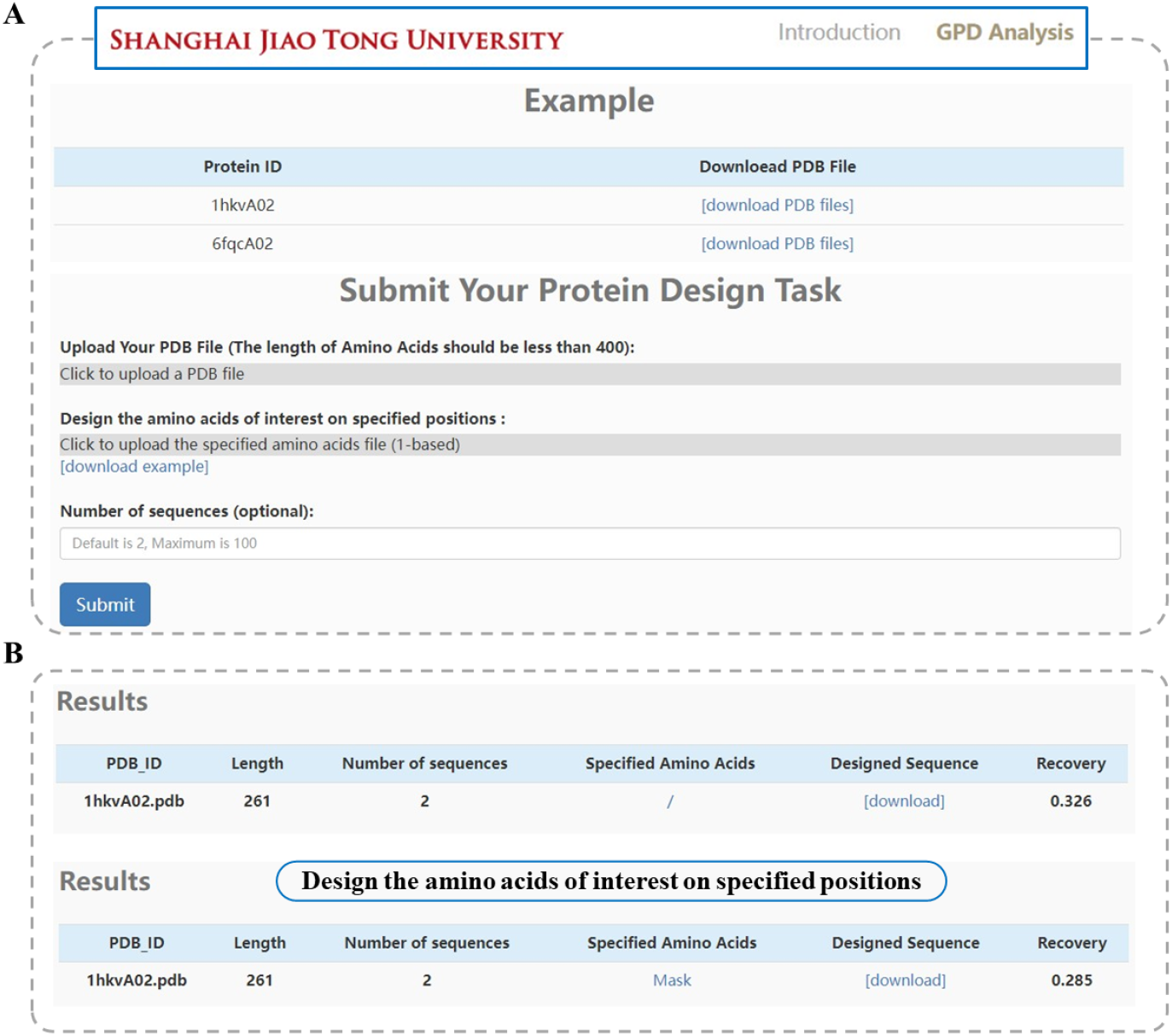
The online tool for GPDGenerator. **a**, The “Analysis” interface of the GPD web server. The user needs to input the PDB file and the specified amino acids (design the amino acids of interest at specified positions are optional **b,** the example output of the web server.

